# Melatonin induces endoreduplication through oxidative DNA-damage triggering lateral root formation in onions

**DOI:** 10.1101/2023.07.31.550947

**Authors:** Sukhendu Maity, Rajkumar Guchhait, Kousik Pramanick

## Abstract

Melatonin (Mel) can regulate lateral root formation, but the underlying molecular mechanisms of Mel-induced lateral root formation are indistinct. This study first time reports the potential ability of melatonin to induce endoreduplication, which in turn could play important roles in developmental reprogramming in plants towards lateral root formation. Pursuant to the results, Mel induces the lateral root formation in onions in a dose-dependent manner with the highest root forming potential in the high concentration (50 µM) of Mel. In consistent with the lateral root formation, the ROS generation in this dose was significantly higher than the control and a low dose (5 µM Mel, Mel_1) group. Co-treatment of ascorbic acid (AsA) with Mel in Mel_2 + AsA group can effectively scavenge the Mel_2 induced ROS, which results in a reduced number of lateral root formation in the co-treatment group. The higher levels of H_2_O_2_ and superoxide in Mel_2 further strengthen the previous report on the role of ROS in lateral root formation. An increase in DNA content was also observed in the Mel_2 group consistent with the level of ROS-induced DNA-damage, suggesting that ROS can induce lateral root formation through oxidative DNA-damage stress and resulting endoreduplication. The results of gene expression analysis through qRT-PCR provide supporting evidence that melatonin, in a dose-dependent manner, can arrest cell-cycle, initiating the endoreduplication cycle in response to oxidative DNA-damage. Observed low level of IAA in primary root tip indicates the DNA-damage and cytokinin-dependent inhibition of auxin polar transport, causing localised IAA accumulation in the zone of differentiation due to auxin bio-synthesis, which in turn triggers lateral root formation in this region in corroboration with endoreduplication and ROS.

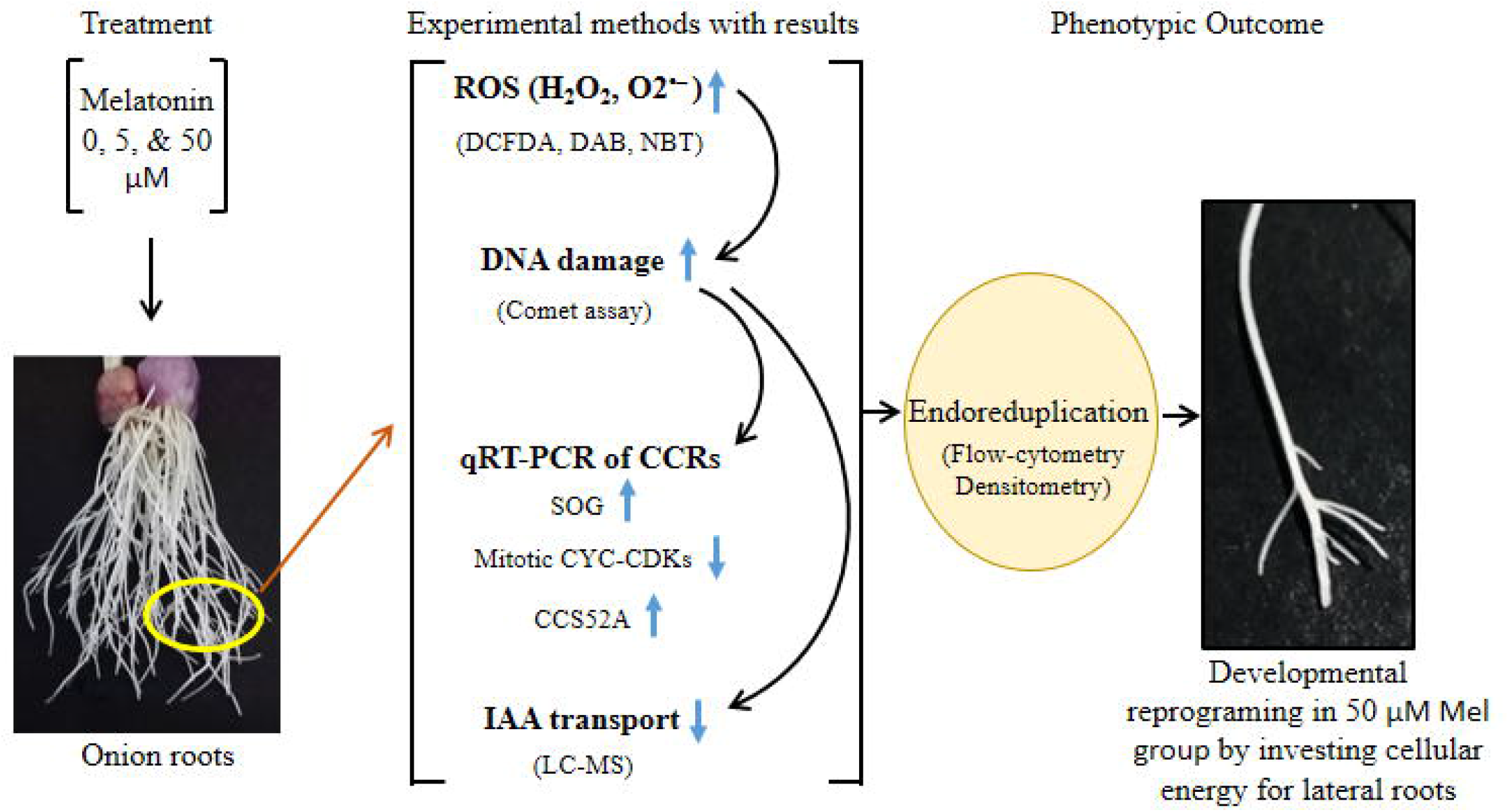

## 1. Introduction

The formation of lateral roots is regulated by hormonal signalling, developmental processes, and environmental cues providing a suitable survival strategy under unfavourable environmental conditions (Chen et al., 2018). During plant organogenesis, meristematic cells partially lose their cell-cycle activity and follow an alternative mode of cell-cycle, called endocycle/endoreduplication/endoreplication, where cells increase their ploidy level through repeated rounds of replication without following mitosis (Kondorosi et al., 2005). The induction of endoreduplication in plants is a physiological response to biotic and abiotic stress such as pathogen attack and DNA-damage, and can also be considered as an adaptive response contributing to the plant ploidy level, metabolism, and survivability/fitness under stressful conditions (De Veylder et al., 2011; Breuer et al., 2014; Kołodziejczyk et al., 2021; Mahapatra and Roy, 2021). Endoreduplication also plays an important role in plant growth, the development of new organs, and cellular differentiation during organogenesis (Cebolla et al., 1999; Shu et al., 2018; Hendrix, 2022). Breuer et al., (2014) reported the positive role of endoreduplication on the branching of *Arabidopsis* leaf trichomes with a prominent over-branching in mutants exhibiting a high level of endoreplication. Thus, the endoreduplication may have important roles in lateral root development that requires experimental validation.

It is also evident that DNA-damage response can arrest cell-cycle and activate endocycles under stressful conditions. DNA-damage in the root tips and leaves of *Arabidopsis* has been proven to induce endoreduplication with a parallel decrease in the expression of mitotic factors (Adachi et al., 2011; Shu et al., 2018). Xu et al., (2016) showed the inhibition of mitotic cyclin-CDKs (cyclin dependent kinase) is a prerequisite for the promotion of endoreduplication. High salinity induces oxidative stress in *Arabidopsis* through elevated ROS production and oxidative damage of DNA, which in turn induces endoreduplication for showcasing a better adaptive response (Mahapatra and Roy, 2021). Regarding lateral root formation, Davis et al., (2016) reported the inhibition of lateral roots in *Arabidopsis* by DNA-damage causing the up-regulation of cytokinin biosynthesis genes. Conversely, De et al., (2022) showed an enhanced level of lateral root formation in *Vigna radiata* after exposure to the low dose of UV-B for a longer period. Two possibilities may come out from these studies: firstly, lateral root development under DNA-damage stress varies with plant species, and secondly, it is the extent of DNA-damage that decides whether lateral root will develop or not.

On the other hand, melatonin (Mel), an endogenous regulator of plant physiology, can serve as a well-known stress alleviator with prominent inducible effects on lateral roots formation (Chen et al., 2018; Chen et al., 2019; Bian et al., 2021). Lateral root formation by melatonin treatment has been reported to be mediated by elevated ROS generation and regulating auxin and cytokinin levels, but both of these pathways may operate independently of each other (Chen et al., 2018; Takahashi et al, 2021; Bian et al., 2021). Therefore, it is also possible that this increased level of ROS by melatonin could lead to oxidative damage of DNA, which requires experimental validation.

In favour of this assumption, it is worth mentioning ribonucleotide reductase (RNR) plays an important role in DNA-damage checkpoint in higher plants, and the expressions of *RNR* and *PCNA1* (proliferating cell nuclear antigen1) have been reported to be up-regulated in melatonin treated *Arabidopsis* (Wang et al., 2017). Proliferating cell nuclear antigen1 (*PCNA1*), histone H2A gene 10 (*HTA10*), and ribonucleotide reductase1 (*RNR1*) are three important S-phase specific genetic markers up-regulated after treatment with melatonin (Wang et al., 2017). These studies may also indicate that melatonin could induce endoreplication.

Melatonin induces lateral root formation by increasing ROS production. Again increasing ROS can lead to oxidative damage of DNA; and as endoreduplication is induced by DNA-damage, it is logical to think about the possible connections among ROS, DNA-damage, endoreduplication, and lateral root formation. Based on the above background studies, it is hypothesised that melatonin induce endoreduplication in a dose-dependent manner by increasing ROS production and DNA-damage stress, and this endoreduplication then redirects the developmental program towards lateral root formation. Therefore the major objective of the present work comprises of the investigation of melatonin-induced DNA-damage and resulting endoreduplication, and finding their interconnections with ROS-signalling and hormonal signalling pathway with regard to the lateral root formation.

## 2. Material and methods

### 2.1. Experimental setup

After collecting the fresh onion bulbs from the local market, dried roots were ablated leaving root initials and the equal size bulbs were set for the treatment. After an initial screening of root forming potential of melatonin (Mel) using different doses (**Fig. S1**), four treatment groups, having three biological replicas in each, were designed with two concentrations of Mel: one low concentration (5 µM Mel) with no root forming potential and one high concentration (50 µM Mel) with root forming potential). Different treatment groups were: Control (no Mel), 5 µM Mel (**Mel_1**), 50 µM Mel (**Mel_2**), and 50 µM Mel + 100 µM ascorbic acid, AsA (**Mel_2 + AsA**). All these experimental groups were left for 12 days in a dark/light cycle of 12 h under laboratory conditions.

### 2.2. Reactive oxygen species (ROS) measurement

For each treatment groups, a total of 3 root tips, one for each biological replica, were stained with 2’,7’-dichlorodihydrofluorescein diacetate, H2DCFDA (Ex/Em: ∼492– 495/517–527 nm) following the protocol of Chakrabarti and Mukherjee (2021). Succinctly, after a brief washing, the cut root tips were incubated for 10 min in the dark at room temperature in 25 µM DCFDA aqueous solution prepared from 4 mM DMSO stock solution. After incubation, samples were thoroughly washed with water and glycerol mounted on clean microscopic slides. Images were captured in AXIO Lab A1 (Zeiss) fluorescence microscope under 10X objective. Corrected total cell fluorescence (CTCF) of DCFDA stained root tips has been estimated as: CTCF = integrated density – (area of selection x mean background fluorescence) using Image-J software.

### 2.3. Histochemical staining of hydrogen peroxide and superoxide ion

Histological staining of H_2_O_2_ and O2^•–^ were performed by staining roots with 1 mg/ml aqueous solution of 3,3’-diaminobenzidine (DAB) and Nitroblue tetrazolium (NBT), respectively, following the protocol of Wang et al., (2023). The pH of DAB was adjusted to 3.8 with concentrated HCl. After an incubation of 6 h for DAB and 30 min for NBT staining, the samples were washed for 1 hr with absolute alcohol to decolourize and then stored at 4 C until observation.

### 2.4. Single cell gel-electrophoresis

For the measurement of ROS induced oxidative damages of DNA, alkaline single cell gel-electrophoresis was performed according to the protocol as standardised in our laboratory (Maity et al, 2023). To briefly state, isolated nuclei in Tris buffer (pH - 7.4) were cast on pre-coated microscopic slides with low melting agarose. After solidification at 4 C, slides were incubated for 20 min in alkaline electrophoresis buffer for denaturation and then the electrophoresis was carried out for 30 min at 250 mA and 25 V in cold room. Slides were neutralised in 400 mM tris (pH - 7.4) with three changes, 10 min incubation in each change, and air-dried. Then, the slides were kept in ice-cold water for 5 min and stained with EtBr (80 µg/ml) for 5 min. Finally, after proper washing, slides were observed in AXIO Lab A1 (Zeiss) fluorescence microscope under 40X objective and 50 comets were scored for each treatment group and analysed in CapsLab (casplab_1.2.3b2.exe) software.

### 2.5. Gene expression analysis through qRT-PCR

The expression of genes, playing crucial roles in DNA replication and endoreduplication, were analysed by quantitative real-time PCR. RNA was extracted from primary root tips using TriZol (Invitrogen) reagent following manufacturers’ instructions. The quality and quantity of isolated RNA were checked in NanoDrop and the integrity was checked in 1.2 % agarose gel. The 1^st^ strand cDNA was synthesised from 1 µg of RNA for each sample using iScript™ cDNA Synthesis Kit (Bio-Rad). This cDNA was then used as the template in qRT-PCR reaction mixture after 10 times dilution. The qRT-PCR reaction mixture was prepared using iTaq SYBR green master mix (Bio-Rad) following the users’ manual and the reaction was run in CFX96 Touch™ Real-Time PCR Detection System (Bio-Rad) under the setup with an initial denaturation at 95 C for 3 min, and 40 cycles of 95 C for 30 sec, 57 – 59 C (annealing temperature) for 10 sec; and melt curve analysis was done using instrument default settings. The C_t_ value of individual genes was normalised to the gene encoding 25s rRNA that showed insignificant or no variation in its expression among different treatment groups. The relative fold change for each gene was calculated in 2^-ΔΔCt^ method. The method of primer design and the list of primers used in this experiment have been given in the supplementary table (**Table - S1**)

### 2.6. Flow cytometry for the quantification of DNA content

Samples were prepared for DNA content analysis by flow cytometry following Loureiro et al., (2006) with slight modifications. Briefly, about 50 mg of primary roots were cut from each experimental group after 4 days of incubation and chopped in 400 µl of ice-cold Tris.MgCl_2_ buffer (200 mM Tris, 4 mM MgCl_2_.6H_2_O, 0·5C% (v/v) Triton X-100, pH 7·5) on ice with a sharp razor to isolate nuclei. The isolated nuclei were then filtered with muslin cloth to remove debris. After a 30 min incubation with 50 µg/ml RNase-A for RNA digestion, the nuclei were stained with 50 µg/ml propidium iodide and then the samples were analysed in BD-Accuri C6 flow cytometer with a slow flow rate and 20,000 events count. From the plot forward scatter vs side scatter, singlets were selected by plotting PI-area vs PI-width and finally, the DNA content was analysed by plotting PI-hight vs event counts in Kaluza, Beckman Coulter, Inc.

### 2.7. Densitometric analysis of DNA content from DAPI stained nuclei

Densitometric analysis of DNA content from DAPI stained nuclei may have an inaccurate estimation of DNA due to the preferential binding of DAPI with AT-rich regions (Noirot et al., 2002). As we aimed to measure the relative DNA content among different experimental groups of the same species, where unless base composition is changed due to treatment, it should not be a major problem. Munyenyembe et al., (2021) measured DNA content through densitometric analysis of DAPI stained nuclei in Arcellinida and Ciliates. Isolated nuclei from root samples as prepared for flow cytometry were also subjected to densitometric analysis of DNA. 10 µl of nuclei isolates were placed on a microscopic slide and air-dried, and then stained with DAPI before image capture in a fluorescence microscope using 40X objective. Ten nuclei were analysed for measuring integrated density in Image-J software.

### 2.8. Orcein staining of chromosome

Root tips were fixed in 1:3 acetic-alcohol for overnight and then hydrolysed in 1N HCL for 10 min at 60 C temperature. The root tips were then stained with 2 % aceto-orcein for 1 hr and finally, the stained root tips were squashed in 45 % acetic acid on grease-free microscopic slides for microscopic observation.

### 2.9. Quantification of auxin (IAA)

For quantitative detection of endogenous auxin using LC-MS, the protocol of Durgbanshi et al., (2005) was followed with slight modification. Briefly, ∼ 200 mg of fresh root tissue from primary root tips and the zone of differentiation, from where lateral roots grow, was ground into the liquid nitrogen and extracted in 1 ml of LC-MS grade methanol:water (80:20 V/V) for 12 h at – 20 °C. Finally, the sample was spinned at 8,000 × g for 10 min. A volume of 5 µl of filtered (PTFE, 0.22 μm, Anpel) supernatant was loaded in Agilent 1200 liquid chromatography having Zorbax Eclipse XDB-C18 (5 lm; 4.6 9 150 mm) column interfaced with Agilent 6410 triple quadrupole spectrometer with ESI ionisation source. Gradient elution was done with water and 0.1% formic acid (solvent A) and methanol with 0.1% formic acid (solvent B) at a constant flow rate of 0.5 ml min^-1^. The gradient was increased linearly from 5% Solvent B to 95% Solvent B over a period of 12 mins and held for 3 mins, after which initial conditions were restored and held for 10 mins. IAA was detected in positive mode with mass of the parent ion at 175.90 m/z and confirmed with the daughter ion at 130.00 m/z. IAA levels were expressed in ng/mg of plant tissue.

### 2.10. Statistical analysis

Data were statistically analysed by one-way ANOVA or nonparametric test (when data was not homogeneously distributed) in IBM SPSS (version 26). Shapiro-wilk normality test was used for testing the distribution of data. When significance was detected, Post Hoc analysis (Tukey test) and Kruskal-Wallis test were performed to compare different experimental group means at 95 % confidence level (p≤0.05). Results were expressed as group mean ± SEM.

## 3. Results

The ability of melatonin for lateral root formation has been reported previously. Out of two different doses of melatonin used in this experiment, only the high dose group - Mel_2 (50 µM Mel) induced lateral roots formation, which was inhibited by ascorbic acid with very reduced numbers of lateral roots in the co-treatment group - Mel_2 + AsA (**Fig. 1A**). This dose-dependent lateral root formation corroborates the study of Liang et al., (2017), where they showed the potential of different doses of melatonin including 50 µM to induce lateral roots, and was also consistent with levels of ROS, DNA-damage, mRNAs of genetic markers, and endoreduplication as discussed later.

**Fig. 1:**
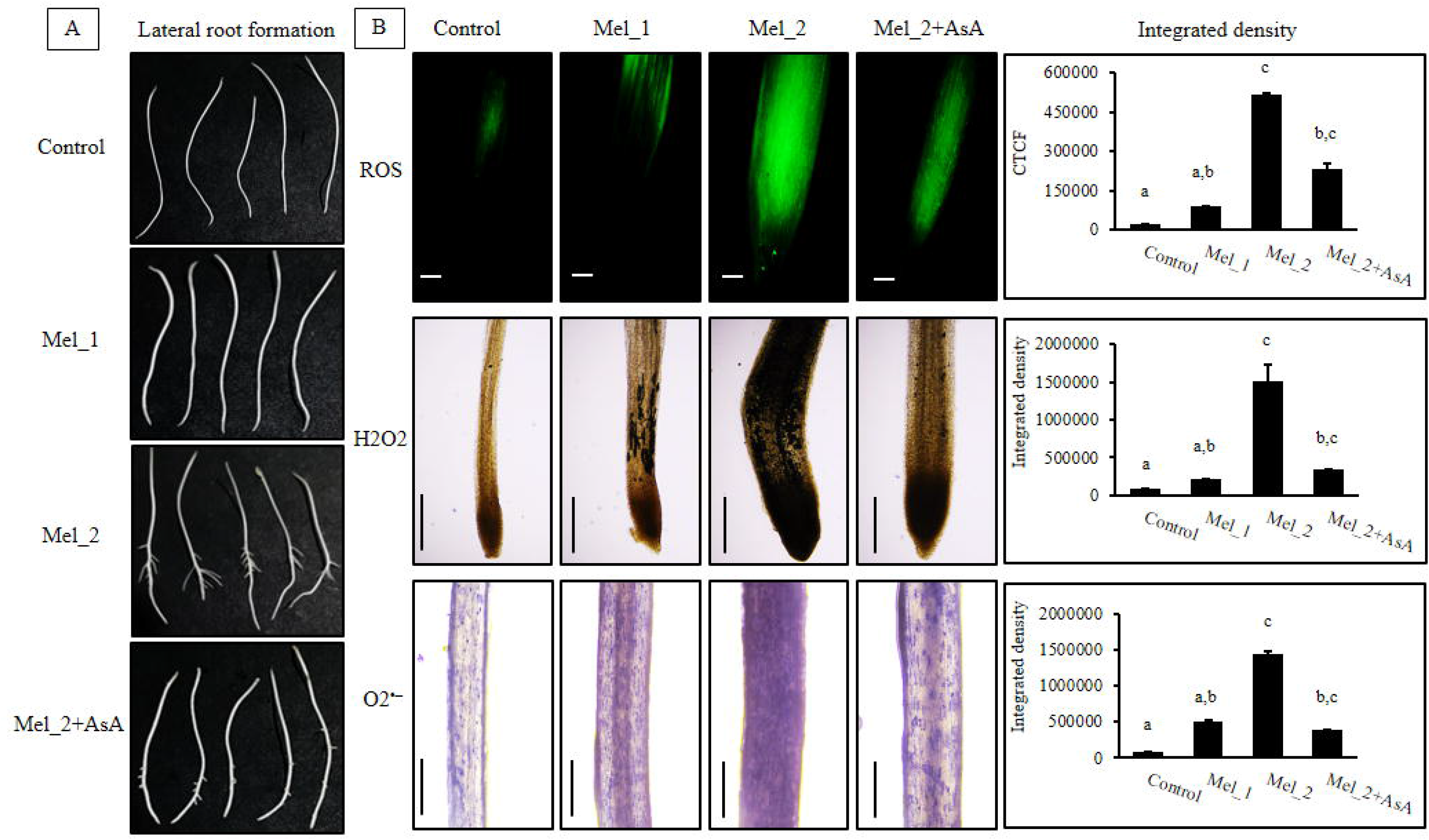
Melatonin (Mel) induced lateral root formation and its regulation by ROS generation in onion. A - Mel induces lateral roots in its high dose group, Mel_2 (50 µM Mel) with no effects on lateral roots formation in the low dose group, Mel_1 (5 µM Mel). The ability of melatonin in inducing lateral roots is remarkably reduced by ascorbic acid treatment in the co-treatment group (Mel_2 + AsA), where 100 µM of ascorbic acid (AsA) was administered along with Mel_2. B - Mel can also trigger redox homeostasis through the up-regulation of ROS, including H_2_O_2_ and superoxide radicals, in a dose-responsive way. The upper row showed the estimation of total ROS in onion roots after DCFDA staining with the highest fluorescence signal for ROS in the Mel_2 group. Bar = 50 µm in upper row images. H_2_O_2_ and superoxide staining with DAB (middle row) and NBT (lower row), respectively also showed similar trend with total ROS, where no significant levels of H_2_O_2_ and superoxide were recorded in the control, Mel_1, and Mel_2 + AsA groups, but not in the Mel_2 group with a significantly high H_2_O_2_ and superoxide levels consistent with the lateral root formation in the Mel_2 group. Bar = 100 µm in microscopic images in middle and lower rows. Different letters indicate significant difference between groups means (p≤0.05).

### 3.1. Dose-dependent ROS production including H_2_O_2_ and O2^•–^ causes oxidative DNA-damage

The H2DCFDA, a non-fluorescent and cell permeable ROS marker, is deacetylated inside cells by cellular esterase and subsequently oxidised by intracellular ROS to highly fluorescent 2’, 7’-dichlorofluorescein (DCF). The measurement of corrected total cell fluorescence (CTCF) after H2DCFDA staining of root samples, showed a significant level of ROS generation in the Mel_2 group as compared to the control. On the other hand, the treatment with ascorbic acid (AsA) debilitates the elevated ROS in the combined treatment group, Mel_2 + AsA (**Fig. 1B**).

The intracellular level of H_2_O_2_ can be detected by histochemical staining with DAB, which is oxidised by hydrogen peroxide in the presence of peroxidase to form dark brown precipitates. The measurement of the integrated density of these brown precipitates in different treatment groups showed a significantly high density in Mel_2, but not in other groups. In Mel_2 + AsA group, the fortification of H_2_O_2_ is impaired by the presence of AsA (**Fig. 1B**).

The histochemical staining with NBT can detect superoxide radicals, where NBT is oxidised by O2^•–^ to generate an insoluble formazan compound. The NBT staining pattern of different treatment groups also showed the same results as observed in DAB staining with a significantly different and highest level of O2^•–^ in the Mel_2 group only, which gets lowered by the presence of ascorbic acid in Mel_2 + AsA group (**Fig. 1B**).

The excess ROS generation in the high dose group suggests the oxidative damage of DNA. Therefore, we assessed the DNA-damage through single cell gel-electrophoresis and the results showed a low level of DNA-damage with no significant difference between control and Mel_1 groups. Whereas, the highest level of DNA-damage was observed in Mel_2 group, which was reduced in the co-treatment group due to ascorbic acid treatment (**Fig. 2**).

**Fig. 2:**
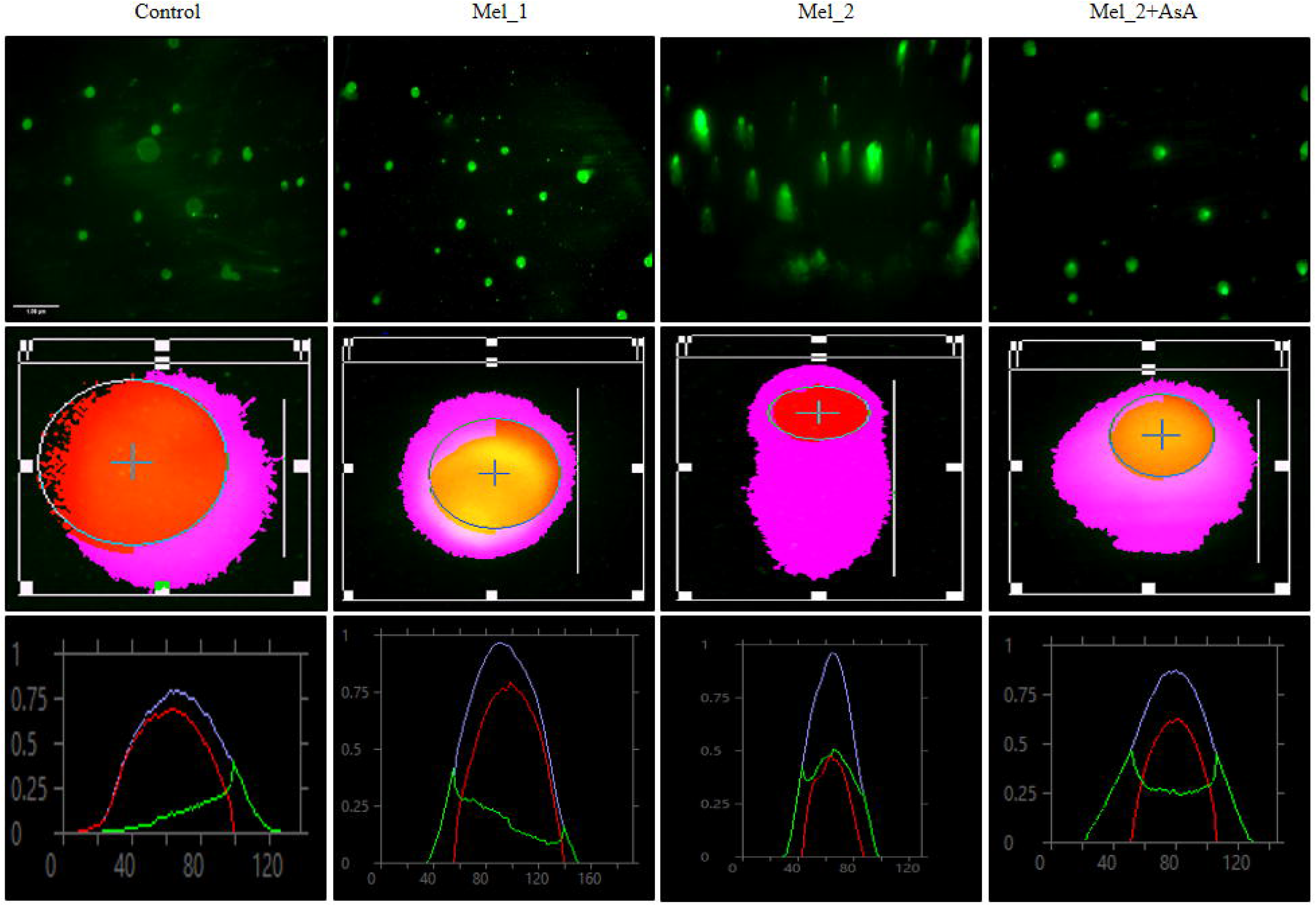
DNA-damage in response to melatonin treatment was measured by comet assay. Melatonin (Mel)-induced ROS formation in different tested doses can be correlated with oxidative damage of DNA. Images in the upper row clearly showed that Mel-induced DNA-damage is dose-dependent with prominent comet formations in the Mel_2 and co-treatment (Mel_2 + AsA) groups. Due to antioxidant activity of AsA, AsA can scavenge Mel_2 induced ROS in the co-treatment group, which in turn debilitates the extent of damage in this group. Bar = 1 µm applied to all microscopic images. Middle and lower rows showed representative image and graph of comet (after analysis in CaspLab) for each treatment groups. In the graphical presentation of comet analysis (lower row images), blue, red, and green colours represent comet, head DNA, and tail DNA, respectively.

### 3.2. DNA-damage in response to melatonin-induced oxidative stress causes cell-cycle arrest and concomitant activation of endocycles

As compared to the control, the high level of DNA-damage as observed in the Mel_2 and co-treatment groups, with tail DNA = 50.36 ± 3.74 % and 25.04 ± 7.47 %, respectively (**Fig. 3**), could lead to the activatation of intracellular signalling cascade leading to cell-cycle arrest and DNA-repair paving the way for stress mitigation and cell survibality. In this study, three sets of genetic markers: DNA-damage responsive markers (*SOG1*, *CYCB*, and *RAD51*), cell-cycle markers (*CDKB*, *CYCA2*, *CYCB*, *CYCD*, and *CDKA*), and endoreduplication specific genetic markers (*CCS52A*, *UBP14,* and *SOG1*) have been analysed through qRT-PCR (**Fig. 3**).

**Fig. 3:**
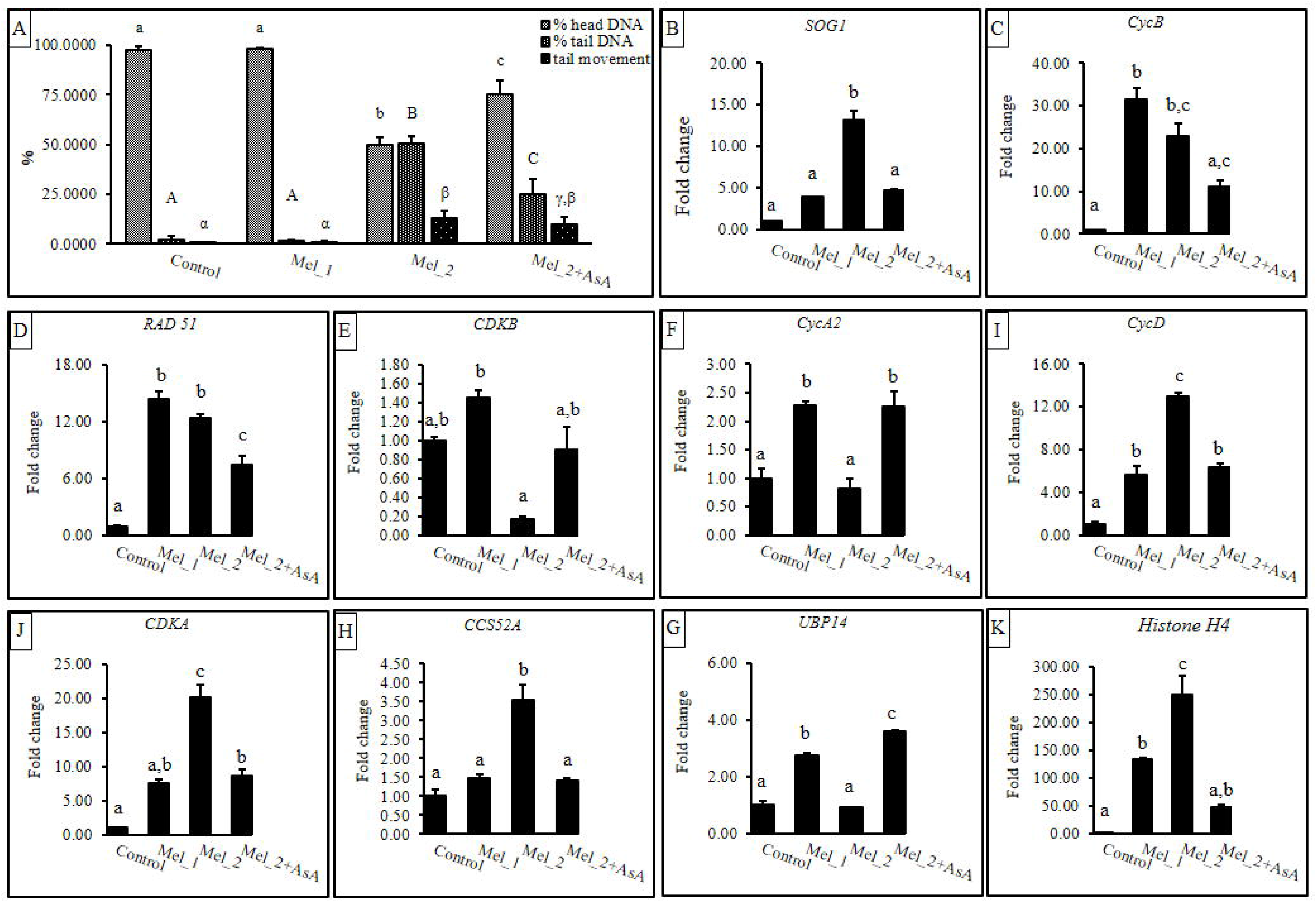
A - Low level of mean head DNA with a parallel increase in tail DNA and tail movement signifies that the oxidative DNA-damage in onion roots after treatment with melatonin was significantly higher in Mel_2 as compared to the control and low dose (Mel_1) group co-treatment groups. Ascorbic acid due to free radical scavenging activity can reduce DNA-damage, as indicated by the significant increase in head DNA and decrease in tail DNA in the co-treatment group (Mel_2 + AsA). B - K, showed the transcript levels of different genetic markers playing crucial role in DNA-repair pathway, cell-cycle regulation, and endoreduplication. The difference in mRNA levels of all of these genetic markers in different treatment groups coincides with the Mel-induced ROS generation, DNA-damage, and the resulting cell-cycle regulation. High level of *SOG1* in Mel_2, as compared to other groups, could signify an efficient operation of DNA repair pathway due to more DNA-damage in this group. SOG1 activates CYCB, which together with CDKB recruits RAD51 leading to DNA repairing process, but significantly low transcript level of *CDKB* in the Mel_2 group can suggest that DNA repairing was not able to mitigate the DNA-damage stress completely, making plants to perceive a stressful condition. As a result, SOG1 turns its way into cell-cycle regulation with significant decrease in mitotic cyclins and CDKs leading to cell-cycle arrest. Significantly high levels of *CCS52* and *UBP14*, as compared to other groups, indicate that cells are arrested with a parallel activation of endoreduplication cycle. The high mRNA levels of S-phase specific markers also support the activation of endocycle in response to DNA-damage in the Mel_2 group. Different letters indicate significant difference between groups means (p≤0.05).

The relative expression level of *SOG1* (suppressor of gamma response 1) was boosted significantly in the Mel_2 group by ∼13.21 folds. As compared to the control, the relative mRNA level of *CYCB* was increased in all treatment groups with a very high level of expressions in Mel_1 (∼ 31.43 folds) and Mel_2 (22.77 folds). *RAD51* expression was higher in both Mel_1 and Mel_2 groups (14.35 and 12.42 folds, respectively) as compared to the control and Mel_2 + AsA groups (∼7.52 folds).

The quantitative assessment of the transcripts of cell-cycle regulating genes can be used to study cell-cycle progression (Gomez et al. 2022). A-type CDKs act throughout the cell-cycle regulating both G1/S and G2/M phase transition while, B-type CDKs only regulates M-phase events in plants (Kondorosi et al., 2005). The expression of mitotic cyclin dependent kinase (*CDKB*) was lower (∼0.18 folds) in the Mel_2 group as compared to other treatment groups. While the expression of mitotic cyclin (*CYCA2*) in Mel_2 (∼0.82 folds) showed a significant difference with Mel_1 (∼2.28 folds) and Mel_2 + AsA (∼2.27 folds) groups, but not with the control group. The mRNA levels of *CYCD* and *CDKA* were higher than control in all three treatment groups: Mel_1, Mel_2, and Mel_2 + AsA with maximum fold change (∼13.00 and 20.19 folds, respectively) in the Mel_2 group. *CCS52A* and *UBP14* showed an opposite pattern of expression with the highest transcript level (∼3.55 folds) of *CCS52A* and the lowest level (∼0.91 folds) of *UBP14* in Mel_2 as compared to other treatment groups. The expression level of the S-phase specific marker, *HISTONE H4* also showed a very high level of expression in the Mel_2 group (∼249.05 folds) as compared to other groups, supporting the ability of melatonin to induce endoreduplication.

### 3.3. Melatonin treatment increases DNA content through endoreduplication in a dose-responsive manner

Flow cytometry analysis of nuclei from different samples revealed that melatonin treatment can increase the DNA content in a dose-dependent manner. In this experiment, 8C and 16C were detected within acceptable limits (∼ 6 %) of the coefficient of variation, CV (Darzynkiewicz et al., 2010). An increase in DNA content was observed in Mel_2 (50 µM Melatonin) group with a gate % of 8C = 1.61 %, and a corresponding CV value of 3.64. There was no significant difference in DNA content among other treatment groups (**Fig. 4A**). To validate increased DNA content resulting from endoreplication in S-phase arrested cells, DNA content was analysed from another treatment group, 50 µM Mel + 40 mM hydroxyurea, HU (**Mel_2 + HU**), where HU inhibits DNA replication by inhibiting ribonucleotide reductase. In this treatment group, the level of 8C (0.46 %) was not significantly different from the control group suggesting the potential ability of Mel to increase DNA content through endoreduplication (**Fig. 4A**).

**Fig. 4:**
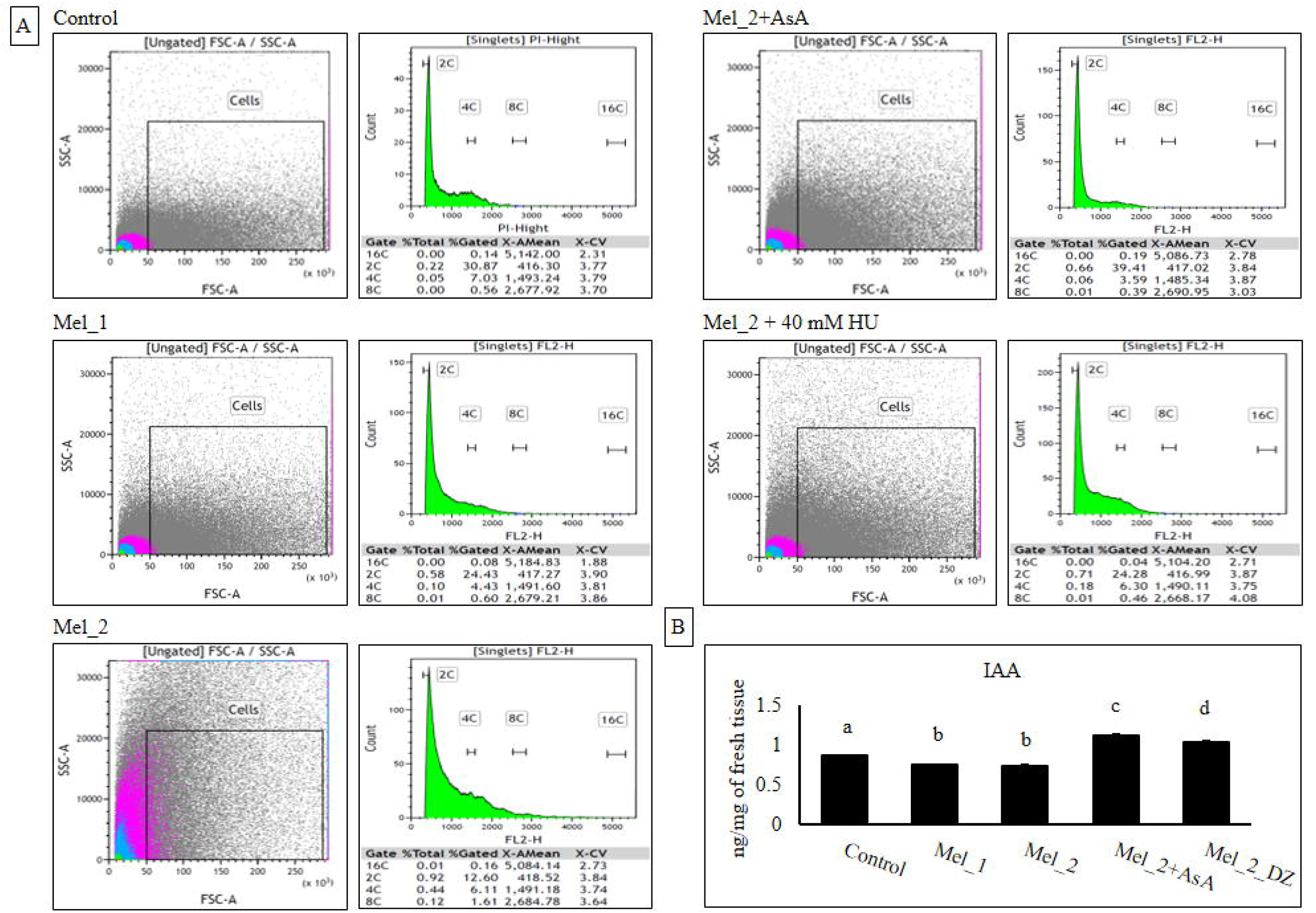
A – Flow cytometric analysis of DNA content in the isolated nuclei of primary root tip samples of different treatment groups after staining with propidium iodide. High level of fluorescence signal (PI-hight) was recorded only in the Mel_2 group as compared to others, indicating an increase in DNA content due to endoreduplication in this group. Melatonin-induced ROS generation and resulting DNA-damage were low in the low dose Mel group (Mel_1) and the co-treatment group (Mel_2 + AsA). As a result, the damaged-DNA was successfully repaired in the Mel_1 and Mel_2 + AsA groups, setting no or less stress, respectively. This endoreduplication was induced after cell-cycle arrest in response to melatonin-induced ROS generation and DNA-damage, as indicated by decrease in DNA content after treatment with hydroxyurea in Mel_2 + HU group, where HU inhibits ribonucleotide reductase and suppresses replication. B – the low level of IAA in primary root tips and high level in the zone of differentiation in Mel_2 group indicates that melatonin reduces the polar trandport of auxin to the primary root tips.

If DNA content is increased, the fluorescence intensity after staining with a DNA-binding fluorochrome will also be increased as compared to the control. The fluorescence intensity of DAPI stained isolated nuclei in Mel_2 was significantly higher than the control and other treatment groups (**Fig. S2**). During endoreduplication, cells are arrested in S-phase, but this cell-cycle arrest would not be complete, therefore, some endoreduplicated cells may pass into the M-phase, which could lead to polyploidy. The presence of some nuclei in the Mel_2 group with chromosome number 32 and 64 as compared to the normal diploid nuclei (2n = 16) strongly suggests the presence of 8C and 16C nuclei, respectively, and Mel treatment can induce endoreduplication (**Fig. S3**).

### 3.4. ROS-induced DNA-damage and endoreduplication together with IAA can modulate lateral root formation by melatonin

In the context of lateral root formation, Mel_1 (5 µM Mel) showed no lateral root formation, whereas, Mel_2 (50 µM Mel) showed the highest lateral root forming potential, which was hampered by ascorbic acid treatment (**Fig.1A**). Results showed that melatonin-induced ROS makes DNA-damage stress leading to cell-cycle arrest and endoreduplication. To examine whether the endoreduplication takes part in lateral root formation or not, a separate experimental setup has been designed, where two experimental groups: Mel_2 and Mel_2 + AsA were incubated for 12 days (**Fig. S4A** and **Fig. S4B**, respectively) and another two experimental groups were designed after an initial incubation of 4 days in Mel_2, and then further incubated for next 8 days in H_2_O-only (**Fig. S4C**) and Mel_2 + AsA (**Fig. S4D**). This study showed melatonin can induce lateral root formation in onions in Mel_2 after 12 days of incubation (**Fig. S4A**). We also observed that after an initial incubation in Mel_2 for 4 days, when we further incubated the samples for next 8 days in H_2_O only (**Fig. S4C**) and in Mel_2 + AsA (**Fig. S4D**), this lateral root formation was not impaired but their growth was reduced, suggesting that the endoreduplication is required for lateral root formation and melatonin-induced ROS is only required in setting the endoreduplication during first 4 days incubation in Mel_2. It is further validated by the absence or presence of very few lateral roots in samples treated for 12 days in Mel_2 + AsA (**Fig. S4B**), where Mel_2 induced ROS is scavenged by AsA from the very beginning of incubation reducing the chance of ROS-mediated endoreduplication.

The level of IAA, as measured by LC-MS, was low (0.74 ± 0.007 ng/mg) in the meristematic region of primary roots, but higher (1.04 ± 0.013 ng/mg) in the zone of lateral root formation in the Mel_2 group. The IAA level in primary root tips was increased (1.13 ± 0.016 ng/mg) in co-treatment group as compared to the control (0.86 ± 0.001 ng/mg) and Mel_2 groups, due to the presence of ascorbic acid (**Fig. 4B; Table - S2**).

## 4. Discussion

### 4.1. Dose-dependent ROS production including H_2_O_2_ and O2^•–^ by melatonin causes oxidative DNA-damage in high dose group

In this study, the potential role of melatonin in inducing endoreduplication and its role in lateral root formation has been investigated in onion using two different doses of melatonin. Phyto-melatonin plays important physiological and developmental roles in response to environmental cues by regulating redox homeostasis through its antioxidant activity and the stimulation of antioxidant enzymes (Khanna et al., 2023). We observed the dose-dependent surge of ROS in onion roots implying the ability of melatonin in setting up a stressful impact on plants in high doses. The high CTCF value of DCF in the Mel_2 group indicates high ROS generation as compared to the low dose (Mel_1) and control groups. In the co-treatment group (Mel_2 + AsA), ascorbic acid can effectively scavenge Mel_2 induced ROS. Ascorbic acid (AsA) serves as an antioxidant and can regulate different physiological and biological responses in plants (Akram et al., 2017; Wang and Huang, 2019). It is also evidenced that AsA mediates cell-cycle progression through prolyl-hydroxylation of required proteins and root elongation in onions (Innocenti et al., 1990; Tank et al., 2017). The levels of H_2_O_2_ and O2^•–^ showed the same pattern with DCF fluorescence establishing the fact that a high dose of melatonin can stimulate ROS production, which is scavenged by AsA in the co-treatment group. This exaggerated ROS generation by melatonin could lead to oxidative DNA-damage. The results of the DNA-damage assay were consistent with the results of ROS with a significantly high level of damage in the Mel_2 group, which is debilitated by ascorbic acid treatment in the co-treatment group (Mel_2 + AsA).

### 4.2. DNA-damage in response to melatonin-induced oxidative stress causes cell-cycle arrest and concomitant activation of endocycles

If the high dose (Mel_2) of melatonin initiates DNA-damage stress, cell-cycle progression would be compromised with concurrent activation of DNA-damage repair pathway. It is reported that DNA-damage response leads to the phosphorylation and activation of SOG1 (PedrozaCGarcia et al., 2022). Activated SOG1, a functional analogue of p^53^, is involved in transcriptionally activating the genes required for DNA repair such as - *CYCB*, which along with CDKB recruits and phosphorylates RAD51 in DNA-damage sites driving homologous recombination (Bourbousse et al., 2018; PedrozaCGarcia et al., 2022). As compared to the control group, a high level of *SOG1* mRNA in all treatment groups in corroboration with those of *CYCB* and *RAD51* indicates the activation of the DNA repair pathway, but the low transcript level of *CDKB* (a B-type CDKs) in Mel_2 suggests that plant could receive DNA-damage stress in this particular dose of melatonin due to unsuccessful repairing. This low CDKB in Mel_2 can also inhibit the cell-cycle progression, as it is reported that in plants, B-type CDKs and A-type cyclins represent mitotic cyclin-CDKs required for M-phase progression (Xu et al., 2016).

On the other hand, the fundamental steps of endocycle induction in plants involve the inhibition of mitotic cycles, therefore, the down-regulation of M-phase specific cyclin-dependent kinases (B-type CDKs) and their corresponding cyclins (A-type cyclins) play a crucial role in shifting cell-cycle to endocycles (Xu et al., 2016). It is also reported that up-regulated CYCA2-CDKB complex is also involved in endocycles inhibition, and the activity of CDKB in this complex depends on the stability of CYCA2 (Chevalier et al., 2011). Low transcript levels of *CDKB* and *CYCA2* in Mel_2 as observed in this experiment thus validate that melatonin arrests cell-cycle with concomitant induction of endoreduplication in response to DNA-damage. CYCA2 is reportedly involved in setting DNA content in plants with the observation that the endoreduplication in *Arabidopsis* leaves is restrained in *CYCA2* overexpression line (Wang et al., 2017).

We also observed elevated levels of mRNA for *CYCD* and *CDKA* in the Mel_2 group. Activated CDKA-CYCD complex is required for the commitment of DNA replication in the S-phase (Chevalier et al., 2011). CDKA-CYCD complex through hyper-phosphorylation of retinoblastoma protein releases E2F/DP, regulating the S-phase specific gene expressions (Himanen et al., 2002). The up-regulation of these genetic markers in Mel_2 thus indicates the increased-level of replication, supporting the occurence of endoreduplication in this treatment group.

Therefore, we checked the mRNA levels of endoreduplication specific genetic markers with the observation that *UBP14* was down-regulated and *CCS52A* was significantly up-regulated in Mel_2. Ubiquitin-Specific Protease-14 (UBP14) and Cell-cycle Switch-52A (CCS52A) act antagonistically with each other serving as inhibitor and activator of APC/C complex, respectively, to drive endoreduplication (Vanstraelen et al., 2009; Xu et al., 2016). UBP14 along with UV-B Insensitive 4 (UVI-4) prevents APC/C complex from mediating proteasomal degradation of CYCA2, which results in the inhibition of endoreduplication. Thus, a low level of *UBP14* and a significantly high transcript level of *CCS52A*, as observed in our results, suggest that UBP14 cannot prevent CCS52A mediated activation of APC/C for proteasomal degradation of CYCA2, ensuring both mitotic exit and the onset of endoreduplication. In this context, it is noteworthy that *De* Veylder et al., (2011) reported the decrease and increase in ploidy levels in plants in CCS52A1 loss-of-function and gain-of-function mutants, respectively. Finally, our flow cytometric observation of 8C DNA content, polyploidy cells, and a significantly high fluorescence intensity of DAPI stained nuclei in the Mel_2 group further support ROS-induced DNA damage and resulting endoreduplication induction.

### 4.3. ROS-induced DNA-damage and endoreduplication together with IAA can modulate lateral root formation by melatonin

A root can be categorized as meristematic zone, transition zone, elongation zone, and differentiation zone starting from root tip to base; and lateral root positioning is restricted in the zone of transition and elongation, whereas, the initiation mostly occurs differentiation zone (Du and Scheres, 2018). It is already evident that Mel can induce lateral roots in plants with different effective concentrations (Ren et al., 2019) through ROS signalling, where H_2_O_2_ serves as a signalling molecule to activate down-stream ROS/Ca^2+^ signalling pathway (Chen et al., 2019; Bian et al., 2021). In this study, we have shown the connecting link between ROS-induced DNA-damage, endoreduplication and lateral root formation. De et al., (2022) reported that the treatment of *Vigna radiate* with a low dose (4.0 kJ m^−2^) of UV-B is coupled with a retarded primary root growth and an increased number of lateral roots, whereas, reduced primary root growth and lateral root numbers had been observed in high dose (8.0 kJ m^−2^) group. In the same study, a significant increase in DNA-damage had been observed in both treatment groups as compared to the control, but with prominent defects in the repair pathway in the case of the high dose UV-B group only.

Therefore, based on De et al., (2022) report and our observation of melatonin-induced DNA-damage and lateral root formation in the same treatment group (Mel_2) and the subsequent reduction of lateral root growth by ascorbic acid treatment in the co-treatment group, we propose that the low level of DNA-damage, initiating DNA damage stress, can regulate lateral root formation in plants.

Now we have answered the question that how DNA-damage stress modulates lateral root formation. The biosynthesis of auxin and their polar transport regulate root meristem size and root growth as reported by Wang et al., (2016), where they observed significantly low expressions of genes for auxin biosynthesis and transport in 600 µM melatonin treated roots with a significant reduction of root meristem size. In the same study, the growth of lateral roots was significantly impaired in this high dose group, but not in low dose groups of melatonin such as: 10 and 100 µM melatonin. Here also, we observed a low level of auxin in primary root tips of 50 µM melatonin (Mel_2) treated samples. On the other hand, Shi et al., (2015) showed that the endogenous auxin level is not affected by treatment with low concentrations of melatonin (10 – 50 µM). Therefore, in our study, the observed low auxin in primary root tips in Mel_2 group is not likely to be mediated directly by melatonin, rather would be an indirect effect, because the melatonin concentration in Mel_2 group is unlikely to impact the auxin bio-synthesis. Zhang et al., (2022) showed that high and low level of melatonin have just opposite effects on IAA level i.e. high melatonin reduces IAA, while low melatonin increases IAA level.

The response of auxin related genes during lateral root formation by melatonin varies with plant species (Chen et al., 2019), indicating the differential role of auxin in melatonin-induced lateral root development. Although, the genome wide analysis in rice has shown melatonin mediates lateral root formation through auxin signalling (Liang et al., 2017). In this context, it is worth mentioning that the overexpression of melatonin biosynthesis gene, N-acetylserotonin-O-methyltransferase, in *Arabidopsis* increases the lateral root numbers in association with a reduction of indole acetic acid (IAA) in roots, but not providing the direct relationship between lateral root formation and IAA level (Bian et al., 2021). Takahashi et al., (2021) reported that DNA-damage in *Arabidopsis* can induce the accumulation of cytokinin in root tip cells with a simultaneous decrease in PIN-FORMED proteins, involved in auxin efflux, leading to a decrease in auxin level in root tips and cell-cycle arrest. Our observations of dose-dependent DNA-damage and reduction of IAA content by melatonin signify that DNA-damage by switching on cytokine genes inhibits polar transport of auxin to primary root tips, which results in low auxin concentration in primary root tips, but a possible high auxin accumulation in the region of lateral root formation.

The accumulation of auxin can activate respiratory burst oxidase homologue, RBOH (Mangano et al., 2017) in the region of lateral root formation. This RBOH then leads to superoxide (O^2•–^) formation in the apoplast, which in turn forms H_2_O_2_ by SOD. Melatonin also activates polyamine oxidase (PAO) and RBOH producing H_2_O_2_ and superoxide in the apoplast (Chen et al., 2019; Bian et al., 2021). Then, apoplastic peroxidase can convert H_2_O_2_ into hydroxyl ions (Bian et al., 2021; Mangano et al., 2017). Again, the apoplastic-ROS (apo-ROS) causes Ca^2+^ influx into cells affecting the expression of cell-cycle regulators and increasing the cytoplasmic-ROS in a feedback reaction through the up-regulation of RBOH (Mangano et al., 2017) (**Fig. 5**). High auxin concentration in the region of lateral root formation may thus help in setting lateral root primordia. Hydroxyl ions, as induced by melatonin and auxin, and auxin activated peroxidase lead to cell-wall loosening/remodelling and lateral root growth exploiting cellular energy conserved by the induction of endoreduplication.

**Fig. 5:**
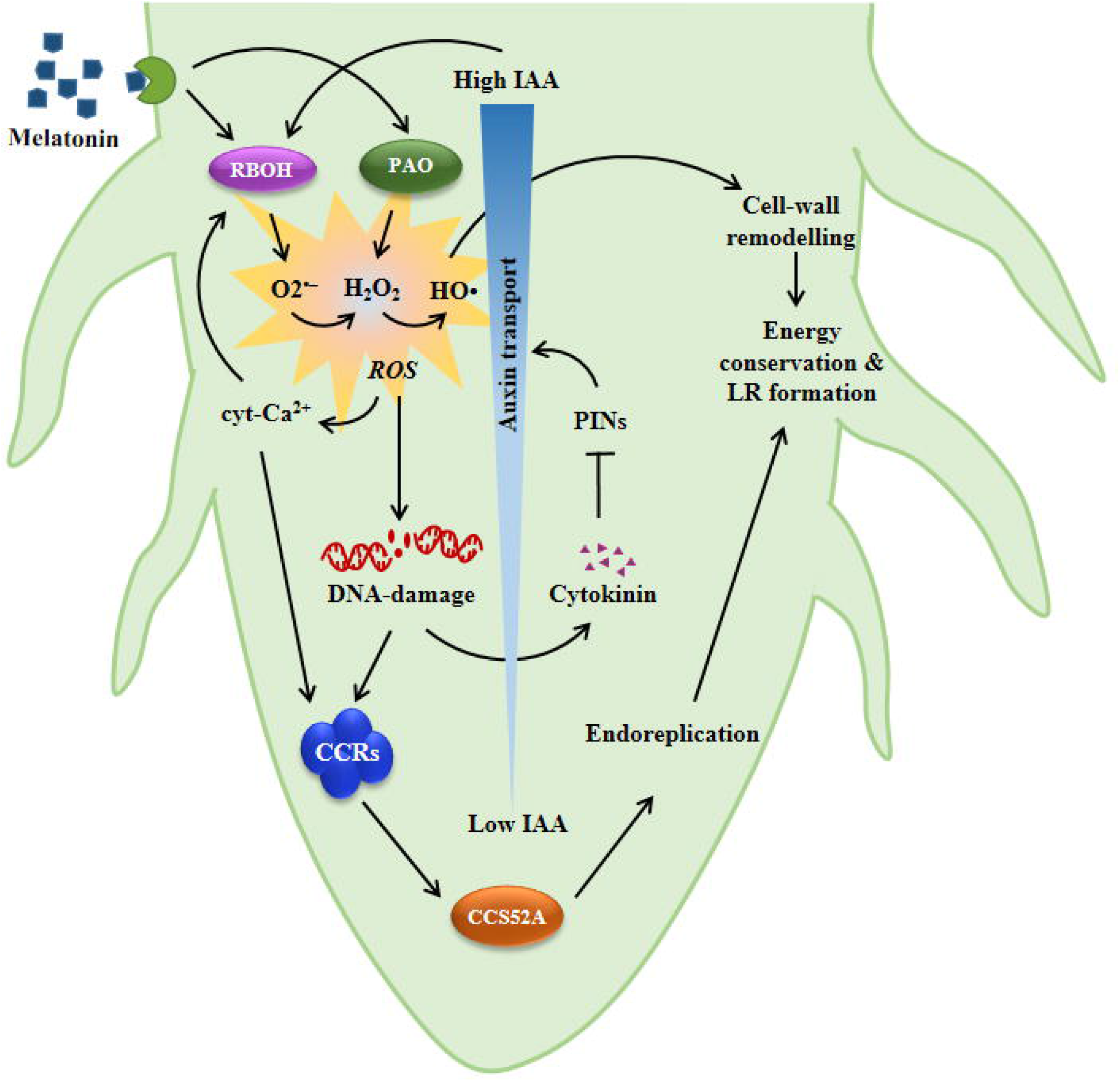
Once melatonin receptors in primary root cells of onion perceive high dose of melatonin it stimulates polyamine oxidase (PAO) membrane bound RBOH (respiratory burst oxidase homologue) to form hydrogen peroxide and superoxide radicals in the apoplast. These free radicals along with peroxidase dependent formation of hydroxyl radicals from hydrogen peroxide initiate an apoplastic oxidative burst. This elevated apoplastic ROS leads to an increased level of Ca^2+^ influx raising intracellular cytosolic Ca^2+^ ions, which in turn in a feedback loop activates RBOH further accelerating ROS generation. This increase in ROS generation could cause oxidative damage of DNA, and therefore, activates DNA-repair pathway for maintaining genomic integrity. In response to DNA-damage, plants can activate *SOG1*, through their transcriptional up-regulation, as a central player in DNA-repair pathway and cell-cycle regulation. SOG1 therefore triggers downstream DNA-repair pathway and also the activity of mitotic cyclins-CDKs by regulating their intracellular levels, leading to cell-cycle arrest. Ca^2+^ dependent signalling pathway can also regulate cell-cycle regulators in response to DNA-damage stress leading to cell-cycle arrest. In this case, melatonin dependent cell-cycle arrest in the high dose group also activates an alternative mode of cell-cycle called endoreduplication through *CCS52A* and *UBP14* up-regulation by SOG1-mediated inhibition of mitotic cyclins-CDKs. Endoreduplication can reportedly increase cellular metabolic activity by increasing ploidy level paving way for survivability under stressful condition. On the other hand, melatonin-induced DNA-damage can elevate the level of cytokinin, which in turn antagonizes the acropetal transport of auxin by inhibiting auxin efflux carrier PINs. Due to this inhibition of auxin transport to the primary root tips, intracellular auxin level can be increased towards to the upper part of primary roots due to localised cellular biosynthesis. This high auxin concentration thus can switch on the reported RBOH mediated ROS generation pathway. High auxin concentration in the region of lateral root formation may thus help in setting lateral root primordia. On the other hand, hydroxyl radicals as induced by melatonin and auxin help in cell-wall remodelling and lateral root growth exploiting cellular energy conserved by the induction of endoreduplication.

## 5. Conclusion

Melatonin-induced lateral root formation in plants is already established with insights into H_2_O_2_ and Ca^2+^ dependent molecular signalling pathway. Here we provide new insights into melatonin-induced lateral root formation connecting the melatonin-induced ROS level, DNA-damage, hormonal signalling, and endoreduplication. Correlation in the occurrence of ROS-accumulation, DNA-damage, endoreduplication, and lateral root formation in high dose (Mel_2) group of melatonin can provide evidence for the ability of melatonin to induce endoreduplication and its modulatory role in lateral root formation. This study also suggests that as long as auxin biosynthesis is not impaired by melatonin, it may act together with auxin rather than acting independently for lateral root induction. Therefore, from this study it is concluded that a low level of endoreduplication is required for melatonin dependent lateral root formation, where melatonin accumulates ROS causing DNA-damage stress, which in turn inhibits auxin transport and arrest cell-cycle with a concomitant onset of endocycle; and eventually, this endocycle reprograms root development towards lateral roots formation so that plants can endure the stressful condition by absorbing more nutrients.

## Acknowledgement

This study is supported by CSIR–JRF grant to the first author (file no. 08/155(0075)/2019-EMR-I) and FRPDF scheme of Presidency University, Kolkata, India. DST-SERB, DBT-BUILDER (BT/INF/22/SP45088/2022), and DST-FIST Govt. of India are also acknowledged in this study for funding research project (EEQ/2018/000275) to the corresponding author and for sponsoring infrastructural facility to DLS (SR/FST/LSI-560/2013(C)). Finally, authors would like to thank Dr.Santanu Chakraborty for allowing us to access Real-Time PCR Detection System (Bio-Rad).

## Authors’ credit

**SM** – Conceptualization, Methodology, Data Curation, Formal Analysis, Software, Writing – Original Draft, **RG** – Visualization, Formal Analysis, Software, **KP** – Conceptualization, Investigation, Funding Acquisition, Supervision, Visualization, Writing – Review & Editing, Validation. All authors have read the paper and approved.

