## Supplementary material for "Melatonin induces endoreduplication through oxidative DNA-damage triggering lateral root formation in onions": supplimentary file

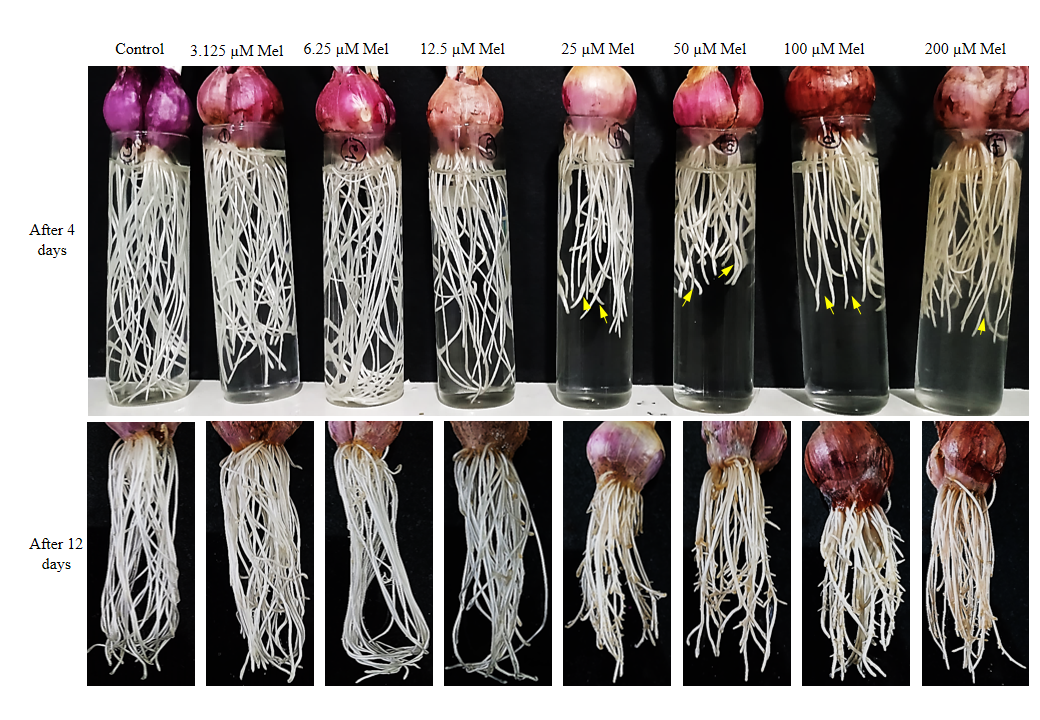

**Fig. S1:** Screening of the ability of melatonin for lateral root formation using different doses. After 4 days melatonin induces endoreduplication in its high dose groups as indicated by swollen root tips pointed by arrow. After 12 days of incubation, melatonin induces lateral roots in high dose groups, but in 200 µM of melatonin group, lateral roots growth is impaired due to inability to combat the stress.

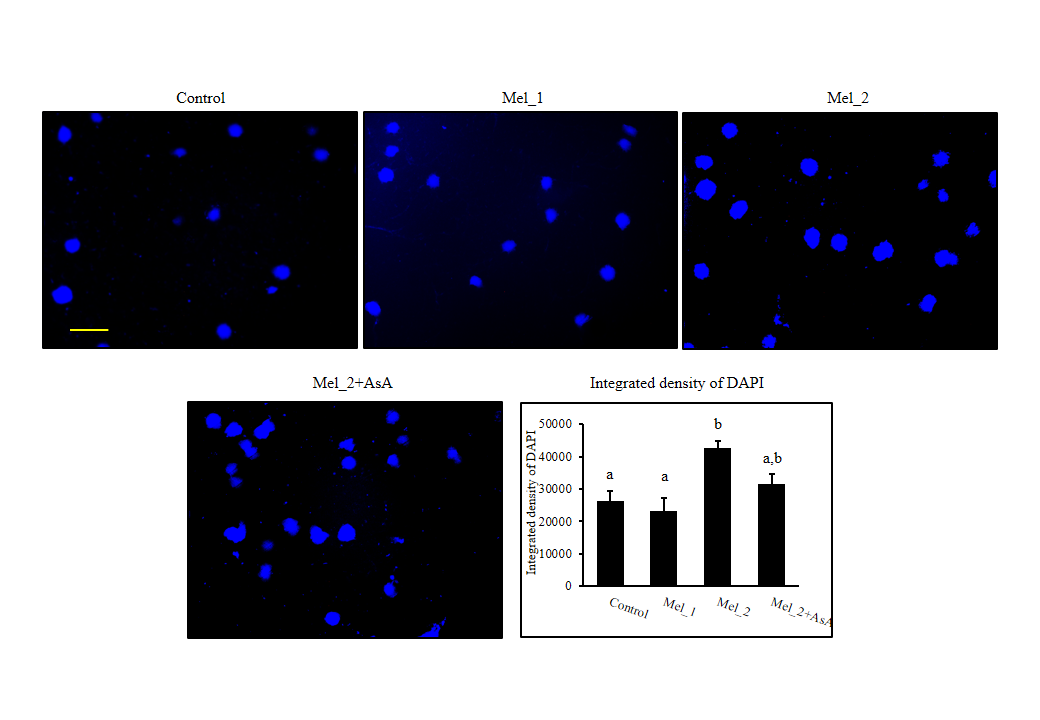

**Fig. S2:** Fluorescence microscopic images of DAPI stained nuclei of different treatment groups, as isolated for Flow-cytometry. The integrated fluorescence intensity analysis showed higher DAPI intensity in the Mel_2 group as compared to other groups. Bar = 100 µm. Different latters indicate significant difference between group means.

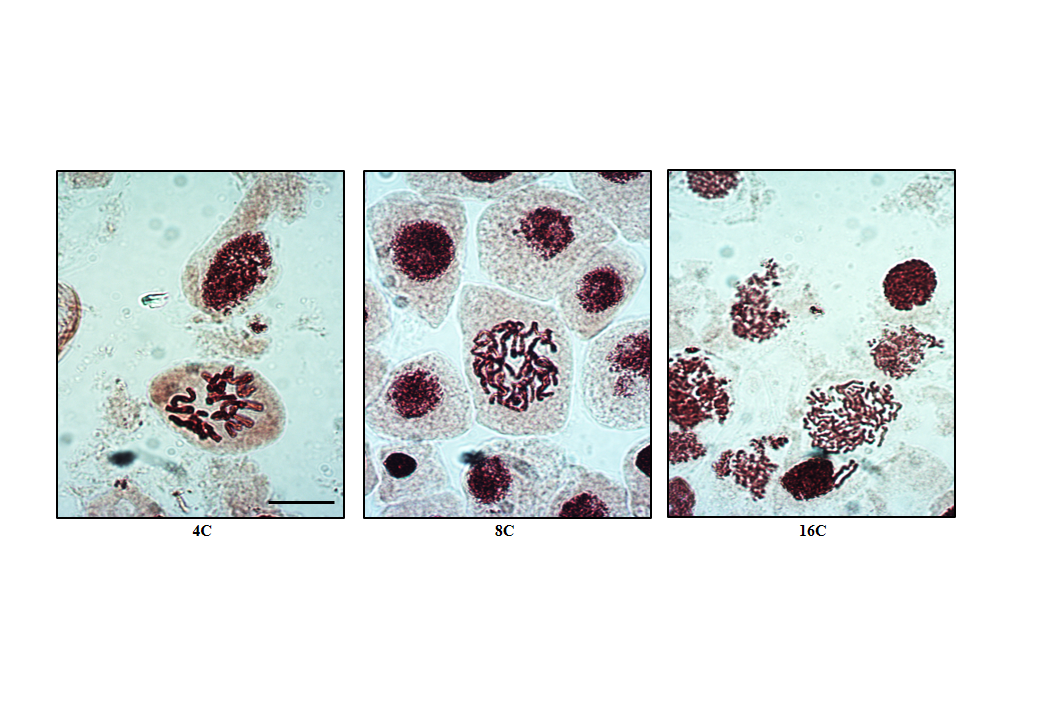

**Fig. S3:** Evidence of polyploidy with DNA content corresponding to 4C, 8C, and 16C. Bar = 0.5 µm applied to all figures.

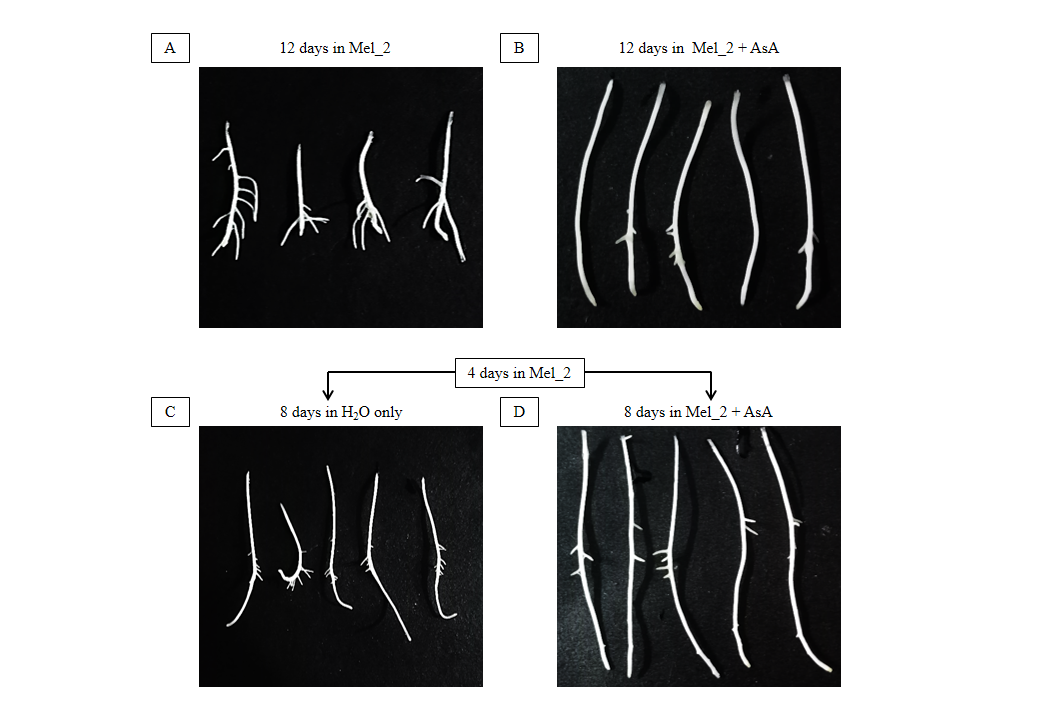

**Fig. S4:** The treatment of onion roots in 50 µM melatonin for 12 days induces lateral root formation, but not in the case of 12 days incubation in Mel_2 + AsA (A, B). In later case, the melatonin-induced ROS is scavenged from the very beginning of incubation, therefore, ROS-induced endoreduplication and the resulting lateral root formation are markedly affected with no or very reduced numbers of lateral roots. To see whether the induction of low level of endoreduplication by melatonin-induced ROS can induce lateral roots, we transferred the samples and further incubated for 8 days in H2O after an initial incubation in the Mel_2 (50 µM melatonin) for 4 days, and observed reduced root growth (C). This result indicates that in the beginning of melatonin incubation for ~ 4 days, ROS is required for lateral root formation, during which, ROS induces endoreduplication by ROS-induced DNA-damage stress; and once after setting off the endoreduplication, ROS is no longer needed for lateral root formation. After an initial incubation in Mel_2, we also incubated for next 8 days in Mel_2 + AsA, and observed similar results as obtained for H2O only i.e. no significant changes in lateral root formation (D). During initial 4 days incubation in the Mel_2, the generated ROS can induce endoreduplication, therefore, after 4 days, scavenging of ROS by ascorbic acid does not impact the lateral root formation, supporting the fundamental role of melatonin-induced ROS in inducing endoreduplication and resulting lateral root formation.

Mel_2_DZ

Mel_2+AsA

Mel_2

Mel_1

Control

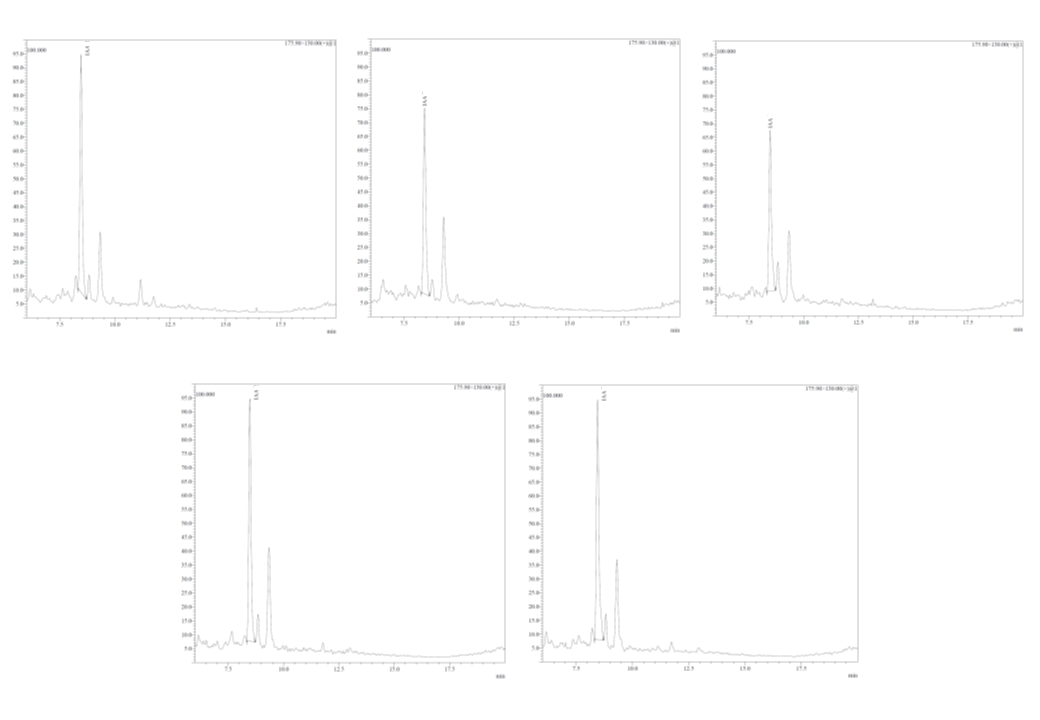

**Fig. S5:** LC-MS Chromatograms of IAA as measured in different treatment groups.

| **Genes** | **Accession no.** | ***Allium cepa* EST accession no.** | **% identity with E-value.** | **Forward and reverse primers (5’ - 3’)** | **Reference** |
| --- | --- | --- | --- | --- | --- |
| *CCS52A2* | NM_117262.3  (*Arabidopsis*) | CF451555.1 | 73.95% (7e-98) | F: CTGGTGTCGTAAAGCGGGAA  R: GTGTTCCTCCTTGTGCCCAT | - |
| *UBP14* | NM_112954.4 (*Arabidopsis*) | [CF434846.1](https://www.ncbi.nlm.nih.gov/nucleotide/CF434846.1?report=genbank&log$=nucltop&blast_rank=1&RID=U32RTSNB016) | 75.43% (6e-110) | F: GTGTTACCGTGCCACCTTCT  R: CCAATTCTTCCTCCCGCAGA | - |
| *Histone H4* | X95689.1  (*Allium cepa*) | - | - | F: GACGCCGTTACCTACACAGA  R: CCCTGACGCTTCAGAGCATA | - |
| *CYCD4* | AJ131636  (*Arabidopsis*) | CF434647.1 | 71.17% (3e-26) | F: CTTGGCTTATCGTTGGCAGC  R: TGGAGTGACTGCGTGGATTC | - |
| *SOG1* | - | CF441566.1 | - | F: TCTGGAACCCGGAAACGTAG  R: TCAACATAGGCTGCTGCTGG | Maity et al., 2023 |
| *RAD51* | - | - | - | F: GCAGCACAGCAGCAAAAGAT  R: ACAGTGCATAGACCTGCGTC | Maity et al., 2023 |
| *CYCA2* | - | - | - | F: CTGGCTTGTTGAGGTTTCCG  R: ACTCTTCGATGGGAGGAGCA | Maity et al., 2023 |
| *CYCB1* | - | - | - | F: TGCACTCTTAGCAGAAGCCC  R: TTCCCTTCTGGTGCAGCAAT | Maity et al., 2023 |
| *CDKA* | - | - | - | F: TTATGGGTACCCCAAACGAA  R: TTGTCCCAAGATCCCTGAAG | Maity et al., 2020 |
| *CDKB* | - | - | - | F: AGGATCACGAGCGAGCATAC  R: AGAGATTGATAGAGATGGACAACAAT | Tank and Thaker, 2014 |

**Table - S1:** List of primers of genetic markers assessed in this study.

| Sample | Retention time (min) | Area | Concentration (ng/mg) ± SEM |
| --- | --- | --- | --- |
| Control | 8.434 | 2,61,134 | 0.86 ± .001 |
| Mel_1 | 8.467 | 2,35,280 | 0.75 ± .002 |
| Mel_2 | 8.477 | 2,24,162 | 0.74 ± .007 |
| Mel_2_DZ | 8.466 | 3,14,921 | 1.04 ± 0.013 |
| Mel_2+AsA | 8.449 | 3,41,194 | 1.13 ± 0.016 |

**Table - S2:** Concentration of melatonin in onion roots during melatonin-induced lateral root formation. DZ = zone of differentiation of primary roots from where lateral roots grow.
